# The Big Five personality traits and CNS arousal in the resting state

**DOI:** 10.1101/2020.10.26.354647

**Authors:** Philippe Jawinski, Sebastian Markett, Christian Sander, Jue Huang, Christine Ulke, Ulrich Hegerl, Tilman Hensch

**Author notes:** **Corresponding author**, Philippe Jawinski, Dr. rer. nat., Department of Psychology, Humboldt-Universität zu Berlin, Unter den Linden 6, 01199 Berlin, Germany.

## Abstract

Based on Eysenck’s pioneering work, CNS arousal has long been considered an encouraging biological candidate that may explain individual differences in human personality. Yet, results from empirical studies remained inconclusive. Notably, the vast majority of published results have been derived from small samples, and EEG alpha power has usually served as exclusive indicator for CNS arousal. In this study, we selected *N* = 468 individuals of the LIFE-Adult cohort and investigated the associations between the Big Five personality traits and CNS arousal by using the low-resolution electromagnetic tomography-based analysis tool VIGALL. Our analyses revealed that subjects who reported higher levels of extraversion and openness to experience, respectively, exhibited lower levels of CNS arousal in the resting state. Bayesian and frequentist analysis results were especially convincing for openness to experience. Among the lower-order personality traits, we obtained strongest evidence for neuroticism facet ‘impulsivity’ and reduced CNS arousal. We regard these findings as well in line with the postulations of Eysenck and Zuckerman and consistent with the assumptions of the ‘arousal regulation model’. Our results also agree with meta-analytically derived effect sizes in the field of individual differences research, highlighting the need for large studies with at least several hundreds of subjects.

## Introduction

Over the past decades, a substantial body of research has focused on the relationship between individual differences in human personality and the underlying biological mechanisms. Aside from a general interest to identify the biological factors that explain the great diversity in human behavior, research in this field has been motivated by theoretical concepts and empirical evidence linking personality traits to mental health outcomes (Maher & Maher, 1994; Strickhouser, Zell, & Krizan, 2017). Beyond this, personality traits have been proposed to constitute vulnerability factors for mental diseases, and affective disorders in particular (Akiskal, Hirschfeld, & Yerevanian, 1983; Barnett et al., 2011; Hensch et al., 2019; Jeronimus, Kotov, Riese, & Ormel, 2016; Klein, Kotov, & Bufferd, 2011). On this account, elucidating the biological basis of personality has not only been argued to provide valuable insights into the etiology of psychiatric diseases, but may also have important implications for identifying at-risk individuals, initiating early preventions, and tailoring treatments.

One of the most prominent trait approaches to describe and measure the structure of human personality is the Five-Factor Model (FFM; Goldberg, 1990; McCrae & Costa, 2008). The FFM is a taxonomy that strives for an economic description of the whole range of individual differences in personality by means of five overarching factors. These ‘Big Five’ personality traits encompass openness to experience, conscientiousness, extraversion, agreeableness, and neuroticism. With some limitations, the five-factor structure of personality has been shown to generalize across languages and cultures, and has been argued to be based on innate biological factors (Macdonald, 1998; McCrae et al., 2000; see also De Raad, 1998). In fact, evidence from twin studies and genome-wide complex trait analyses suggests that a substantial proportion of the Big Five variance is accounted for by genetics (Bouchard & McGue, 2003; Lo et al., 2017; Vernon, Martin, Schermer, & Mackie, 2008). However, the biological mechanisms that bridge the effects of genetic variation on human personality still remain elusive. In order to provide an explanatory biological basis of human personality, various neuropsychological trait theories have been postulated, with Eysenck’s Arousal-Activation Theory of Extraversion and Neuroticism having attracted particular attention (Brocke & Battmann, 1992; Eysenck, 1967).

Eysenck’s Arousal-Activation Theory builds upon the early 1960s’ psychophysiological activation theories, according to which the ascending reticular activation system (ARAS) regulates central nervous system (CNS) arousal (Duffy, 1962; Malmo, 1959). Eysenck distinguishes two components of his conceptual nervous system: the reticulo-cortical brain system (i.e., ARAS) and the reticulo-limbic visceral brain system (VBS; Matthews & Gilliland, 1999). Excitation of the ARAS by incoming stimuli is referred to as ‘arousal’, whereas the excitation of the VBS by emotional stimuli is referred to as ‘activation’. An increase in activation has arousing effects, while arousal may also occur without activation (i.e., a unidirectional relationship). Eysenck postulated that extraverted individuals possess, on average, relatively low habitual levels of CNS arousal in the resting state, which he traces back to a higher ARAS activation threshold (Brocke & Battmann, 1992). As a compensatory mechanism, they engage in arousal-enhancing behavior by seeking human interactions as well as novelty, change and excitement. In comparison, Eysenck describes neurotic individuals as emotionally hypersensitive, which he attributes to a lower activation threshold of the VBS (Brocke & Battmann, 1992). According to Eysenck, individuals with high levels of neuroticism are more susceptible towards stress and show a prolonged autonomic stress response.

The Arousal-Activation Theory has served as theoretical framework in numerous empirical studies (Küssner, 2017; Matthews & Gilliland, 1999). Eysenck himself referred to the alpha range of the human Electroencephalogram (EEG) as the standard measure of CNS arousal (Matthews & Gilliland, 1999). In line with Eysenck’s postulations, a number of studies demonstrated higher resting-state EEG alpha power (indicating lower CNS arousal) in extraverted relative to introverted individuals (Gale, Coles, & Blaydon, 1969; Gale, Edwards, Morris, Moore, & Forrester, 2001; Hagemann et al., 2009; Smith et al., 1995). Several other studies failed to provide supportive evidence (Beauducel, Brocke, & Leue, 2006; Hagemann et al., 1999; Matthews & Amelang, 1993; Schmidtke & Heller, 2004). In addition, some investigators used EEG beta power as CNS arousal indicator and revealed both supporting and opposing evidence (Gale et al., 1969; Gram, Dunn, & Ellis, 2005; Matthews & Amelang, 1993). In sum, empirical investigations addressing the link between extraversion and CNS arousal have provided only inconsistent evidence for Eysenck’s postulations.

A few studies also reported on the relationship between neuroticism and arousal. Based on Eysenck’s postulations, researchers have argued that the habitual level of arousal may tend to be higher in labile (N-) extraverts and introverts when compared to their stable (N+) counterparts (Brocke, Netter, & Hennig, 2004). Consistent with this assumption, investigations in laboratory settings – including resting-state assessments – have previously been shown to elicit an arousal-enhancing ‘first day in lab effect’ (Huang et al., 2015) similar to the ‘first night’ effect in sleep medicine (Hirscher et al., 2015). This may especially affect individuals with high levels of neuroticism, who have been proposed to be more vulnerable towards stress. Notably, enhanced arousal levels in neurotic individuals would also tie in with the substantial genetic overlap demonstrated between neuroticism and major depression (Baselmans et al., 2019; Lo et al., 2017), with the latter having repeatedly been linked to enhanced and ‘hyperstable’ CNS arousal levels in the resting state (Hegerl, Wilk, Olbrich, Schoenknecht, & Sander, 2012; Sander, Schmidt, Mergl, Schmidt, & Hegerl, 2018; Schmidt et al., 2016, 2017; Ulke et al., 2017; Ulke, Tenke, et al., 2019; Ulke, Wittekind, et al., 2019). Despite these converging lines of research, available EEG studies have not yet provided supportive evidence for an association between CNS arousal and neuroticism (Gale et al., 2001; Hagemann et al., 2009, 1999; Savage, 1964). It should be noted, though, that the vast majority of published results on both neuroticism and extraversion have been derived from small samples with fewer than 100 subjects.

In comparison to previous approaches that predominantly used EEG alpha power as exclusive indicator for CNS arousal, above-mentioned studies that demonstrated a link between CNS arousal and depression made use of the Vigilance Algorithm Leipzig (VIGALL), an EEG- and EOG-based analysis tool that utilizes low-resolution electrotomography (Sander, Hensch, Wittekind, Böttger, & Hegerl, 2016). VIGALL is typically applied to fifteen to twenty-minute resting-state recordings and incorporates information on the cortical distribution of the frequency bands alpha, delta, and theta. Beyond this, VIGALL features adaptive procedures that account for the individual differences in alpha peak frequency and EEG total power. Primarily, VIGALL was developed for investigating arousal disturbances in psychiatric samples and to objectively test the assumptions of the ‘arousal regulation model of affective disorders and attention-deficit hyperactivity disorder’ (Hegerl & Hensch, 2014). Similar to Eysenck’s theory, the arousal regulation model postulates that depressive- and manic-like behavior partly reflects an autoregulatory attempt to reduce and enhance habitual high and low arousal levels, respectively. A particular emphasis is put on the regulation of arousal, which is postulated to be unstable in clinical syndromes such as ADHD and mania and is expressed, at the behavioral level, in hyperactivity and sensation seeking (similar to the behavior frequently observed in overtired children). Major depression, in contrast, is postulated to be characterized by enhanced and hyperstable arousal, which is behaviorally expressed in avoidance of additional external stimulation. Noteworthy, by applying VIGALL, a number of empirical studies addressing arousal in depressive, bipolar, and ADHD patients have provided supportive evidence for the assumptions of the arousal regulation model (Hegerl et al., 2012; Strauß et al., 2018; Ulke et al., 2017; Ulke, Wittekind, et al., 2019; Wittekind et al., 2016). In addition, VIGALL has been validated in an fMRI and PET study (Guenther et al., 2011; Olbrich et al., 2009), against evoked potentials and parameters of the autonomous nervous system (Huang et al., 2017, 2018; Olbrich et al., 2011), against the Multiple Sleep Latency Test (Olbrich et al., 2015), and in a large study addressing the agreement with subjective ratings (Jawinski et al., 2017). These previous encouraging results raise the question, whether the application of VIGALL may leverage investigations on the role of arousal in human personality.

Against this background, we here sought to examine the relationship between the Big Five personality traits and CNS arousal in the resting state by making use of the EEG- and EOG-based analysis tool VIGALL. In accordance with previous theoretical and empirical indications, we hypothesized that CNS arousal is negatively associated with the personality trait extraversion and positively associated with neuroticism. Notably, each Big Five personality trait has been demonstrated to genetically overlap with psychiatric disorders (Lo et al., 2017), and each of the respective psychiatric disorders has been proposed to possess arousal-related pathophysiologies (Hegerl & Hensch, 2014). On this account, we here examined the potential associations between CNS arousal and each Big Five personality trait. Given the relatively weak effect sizes in personality and individual differences research (Gignac & Szodorai, 2016; Schäfer & Schwarz, 2019), we considered a sample of several hundreds of participants to derive our estimates. In this vein, we sought to contribute empirical evidence to the so-far unresolved issue of whether basic personality dimensions are reflected in habitual levels of arousal.

## Methods and Materials

In the following sections, we report how we determined our sample size, all data exclusions (if any), all manipulations, and all measures in the study (Simmons, Nelson, & Simonsohn, 2012). All analysis scripts have been made publicly available on the repository of the Open Science Framework (https://doi.org/10.17605/osf.io/ud38w). The original data will be accessible via the Leipzig Health Atlas (https://www.health-atlas.de) upon publication of the peer-reviewed article.

### Sample

Participants were drawn from the LIFE-Adult study, a population-based cohort study of 10,000 inhabitants of the city of Leipzig, Germany (Loeffler et al., 2015). The scope of LIFE-Adult is to examine prevalences, genetic predispositions, and lifestyle factors of civilization diseases. All subjects underwent a comprehensive medical assessment program and completed various psychological surveys. We considered subjects with available resting-state EEG and NEO Personality Inventory data (562 subjects aged 40-79 years). Of these, we selected subjects who reported no current intake of EEG-affecting drugs and had no prior diagnosis of stroke, multiple sclerosis, Parkinson’s disease, epilepsy, skull fracture, cerebral tumor, or meningitis (leaving 533 subjects). Based on a structured clinical interview for DSM-IV axis I disorders, we selected subjects without a history of psychotic disorders or substance dependence, and who were free of current anxiety and affective disorders (leaving 528 subjects). Moreover, EEGs with substantial artifacts (≥ 15% of all EEG segments) and those showing low-voltage alpha, alpha variant rhythms, or pathological activity were not included. This resulted in *N* = 468 eligible subjects (246 females; mean age: 58.5 years). Participants gave written informed consent and received an expense allowance. All procedures were conducted according to the Declaration of Helsinki and were approved by the Ethics Committee of the University of Leipzig (263-2009-14122009).

### Questionnaire

Subjects completed the German version of the revised NEO Personality Inventory (NEO-PI-R; Costa & McCrae, 1992; Ostendorf & Angleitner, 2004). The NEO-PI-R is a widely used self-report questionnaire that enables measuring the personality traits neuroticism, extraversion, openness to experience, agreeableness, and conscientiousness. The NEO-PI-R consists of 240 items and ratings are made on a five-point scale ranging from ‘strongly disagree’ to ‘strongly agree’. Item scores are aggregated to the five NEO personality dimensions. The internal consistency (Cronbach’s alpha) of the five overarching factors has been reported to range from 0.87 to 0.92 (Ostendorf & Angleitner, 2004). Test-retest reliability (1-month interval) has been reported to range from 0.88 to 0.91. Further, the NEO-PI-R allows to calculate scores for thirty personality facets, six facets per factor. The internal consistency and test-retest reliability of the facets has been reported to range from 0.53 to 0.85 and 0.48 to 0.90, respectively (Ostendorf & Angleitner, 2004). NEO personality dimension and facet scores were transformed into sex- and age-normalized T-scores according to the NEO-PI-R manual.

### Physiological data collection and processing

Physiological data collection and processing was carried out as previously described (Jawinski et al., 2019, 2015). EEG assessments were conducted according to a standardized operating procedure. Assessments took place at three time slots: 8:30 am, 11:00 am, and 1:30 pm. During the twenty-minute resting condition, subjects lay on a lounge chair within a light-dimmed sound-attenuated booth. Subjects were instructed to close their eyes, relax and not to fight any potential drowsiness. In order to achieve similar initial levels of arousal activation, all subjects completed a brief arithmetic task immediately before the onset of recording. EEGs were derived from 31 electrode positions according to the extended international 10-20 system. Two bipolar electrodes served to record vertical and horizontal eye movements (EOGs). EEGs were recorded against common average reference with AFz ground. We used a QuickAmp amplifier (Brain Products GmbH, Gilching, Germany) and sampled recordings at 1000 Hz. EEG offline processing was carried out using Brain Vision Analyzer 2.0 (Brain Products GmbH, Gilching, Germany). EEGs were filtered (70 Hz low-pass and 0.5 Hz high-pass with 48 dB/Oct slope, 50 Hz notch) and rectified from eye movement, sweating, cardiac, and muscle artifacts using Independent Component Analysis (ICA). Graph elements (sleep spindles and K-complexes) were manually marked by experienced raters as previously described (Jawinski et al., 2017). Please see the publicly available VIGALL 2.1 manual for further preprocessing details (Hegerl et al., 2016).

### Assessment of brain arousal

The assessment of brain arousal was carried out as described elsewhere (Jawinski et al., 2019, 2017). EEG-vigilance served as indicator for brain arousal and was measured using the Brain Vision Analyzer add-on VIGALL 2.1 (https://www.deutsche-depressionshilfe.de/forschungszentrum/aktuellestudien/vigall-vigilance-algorithm-leipzig-2-1; Hegerl et al., 2016). Based on the cortical distribution and spectral composition of EEG activity, VIGALL assigns one of seven EEG-vigilance stages to each one-second EEG segment. EEG-vigilance stages correspond to active wakefulness (stage 0), relaxed wakefulness (stages A1, A2, A3), drowsiness (stages B1, B2/3), and sleep onset (stage C). Notably, stages A1-3 are characterized by predominant alpha activity, which may indicate relatively enhanced CNS arousal during eyes-closed resting-state conditions where stages of drowsiness (delta- and theta-activity) and sleep onset (occurrence of K-complexes and sleep spindles) are frequently observed. Therefore, the range of arousal stages implicated in the present study extends traditional approaches where higher EEG alpha power (relaxed wakefulness) has been used as exclusive indicator for reduced CNS arousal. We transformed assigned EEG-vigilance stages into values ranging from 7 (active wakefulness) to 1 (sleep onset) and calculated three outcome variables: mean vigilance, stability score, and slope index. Variable ‘mean vigilance’ provides an estimate for the average level of EEG-vigilance during rest. The variables ‘stability score’ and ‘slope index’ particularly focus on the dynamics of EEG-vigilance. Lower scores indicate lower average levels and steeper declines, respectively, of EEG-vigilance. All three outcome variables have been validated, have been found test-retest reliable, and have previously been used as default parameters to summarize complex EEG-vigilance time-courses (Huang et al., 2017, 2015; Jawinski et al., 2017; Jawinski, Mauche, et al., 2016; Jawinski, Tegelkamp, et al., 2016).

### Statistical analyses

The internal consistency of the NEO personality dimensions and facets was calculated using SPSS Statistics 25.0 (IBM corp.; Armonk, New York, USA). All frequentist analyses were carried out using Matlab R2018a (The MathWorks Inc., Natick, Massachusetts, USA). The nominal level of significance was set at *p* < .05 (two-tailed). Further, p-values were adjusted by applying the False Discovery Rate (FDR) procedure according to Benjamini and Hochberg (1995). Associations with FDR < 0.05 were regarded as significant after multiple testing correction. In addition, we sought to derive evidence for the alternate and null hypothesis, respectively, by calculating Bayes factors. Bayes factors reflect the likelihood ratio between the alternate and null hypothesis (BF_10_). Bayesian analyses were conducted with a moderate symmetrical 1/3 beta prior width using package ‘BayesFactor’ (Morey & Rouder, 2018) for R 4.0.1 (R Core Team, 2017).

First, we carried out Spearman correlations between the higher-order NEO personality dimensions (sex- and age-normalized T-scores) and the three EEG-vigilance variables (mean vigilance, stability score, and slope index). Next, we generated a permutation-based quantile-quantile plot (qq-plot) to examine whether the distribution of observed p-values differs from a random p-value distribution under the null hypothesis. On this account, for the set of 15 observed p-values (5 NEO personality dimensions x 3 EEG-vigilance variables), one million sets of 15 expected p-values were derived from correlations after data permutation. Original correlations within the domain of personality traits and the domain of EEG-vigilance variables were preserved, whereas original correlations between these domains were removed through random shuffling. Subsequently, in order to identify facets that particularly contribute to the observed associations, we conducted exploratory Spearman correlations between the thirty NEO personality facets (sex- and age-normalized T-scores) and the three EEG-vigilance variables. By analogy to the higher-order ‘Big Five’ analyses, we also generated a permutation-based qq-plot for the NEO personality facets. Analyses were repeated with sex, age, and daytime of EEG-assessment serving as covariates.

### Statistical power

Power analyses were conducted using R package *pwr* (version 1.3-0; Champely, 2020), with effect sizes quantified as Spearman’s rho (r_S_). Given *N* = 468 and α = .05, power calculations revealed that associations with true effect sizes of r_S_ = 0.052, r_S_ = 0.091, and r_S_ = 0.129 were identified with a chance of 20%, 50% and 80% (1-β), respectively. After Bonferroni-correction (α = .0033; resembling the most conservative case where the FDR procedure ends at the smallest observed p-value), power calculations revealed that associations with true effect sizes of r_S_ = 0.097, r_S_ = 0.135, and r_S_ = 0.173 were identified with a chance of 20%, 50% and 80% (1-β), respectively. Supplementary Figure S1 shows the probabilities of associations to reach the threshold of significance, given true effect sizes of up to r_S_= 0.4.

## Results

The descriptive statistics for the five higher-order NEO personality traits and the three EEG-vigilance variables are shown in Table 1.

**Table 1.** Descriptive statistics of NEO-PI-R scores and VIGALL 2.1 variables of EEG-vigilance

| N = 468 | Cronbach's $\alpha$ | Mean | SD | Q1 | Q2 | Q3 | Min | Max | Skew | Kurt |
| --- | --- | --- | --- | --- | --- | --- | --- | --- | --- | --- |
| **NEO-PI-R** |  |  |  |  |  |  |  |  |  |  |
| Neuroticism | 0.906 | 44.95 | 8.77 | 39.00 | 45.00 | 52.00 | 20.00 | 71.00 | -0.23 | -0.29 |
| Extraversion | 0.899 | 51.69 | 9.83 | 45.00 | 52.00 | 58.00 | 21.00 | 79.00 | -0.02 | 0.18 |
| Openness | 0.868 | 46.85 | 8.45 | 41.00 | 46.00 | 51.00 | 20.00 | 72.00 | 0.32 | 0.32 |
| Conscientiousness | 0.836 | 53.02 | 8.87 | 47.00 | 52.00 | 59.00 | 26.00 | 80.00 | 0.10 | 0.09 |
| Agreeableness | 0.881 | 54.73 | 9.04 | 48.00 | 54.00 | 61.00 | 30.00 | 80.00 | 0.30 | 0.03 |
| **EEG-vigilance** |  |  |  |  |  |  |  |  |  |  |
| Mean vigilance | - | 5.09 | 1.06 | 4.43 | 5.40 | 5.89 | 1.93 | 6.76 | -0.90 | -0.03 |
| Stability score | - | 9.20 | 4.19 | 6.00 | 9.00 | 13.00 | 1.00 | 14.00 | -0.58 | -0.94 |
| Slope index | - | -1.52 | 0.92 | -2.26 | -1.38 | -0.74 | -4.26 | 0.73 | -0.57 | -0.58 |

SD: standard deviation, Q1: quartile 1, Q2: quartile 2 (median), Q3: quartile 3, Min: minimum observed value; Max: maximum observed value; Skew: skewness, Kurt: excess kurtosis

The internal consistency (Cronbach’s alpha) of the five NEO personality dimensions ranged between 0.84 and 0.91 and was thus comparable to previous reports (Ostendorf & Angleitner, 2004). The NEO personality dimensions were significantly intercorrelated (suppl. Table S1), with the strongest correlation observed between extraversion and openness to experience (r_S_ = .477, p = 5E-28). The internal consistency of the NEO personality facets ranged between 0.46 and 0.83 (Cronbach’s alpha and intercorrelations shown in suppl. Figure S2). Intercorrelations between EEG-vigilance variables reached r_S_ ≥ 0.82 (suppl. Table S2). Regarding covariates, we observed that younger participants and those who underwent the EEG assessment at later daytime exhibited a lower EEG-vigilance (e.g. mean vigilance; age: r_S_ = .168, p = 3E-4; daytime: r_S_ = -.155, p = 8E-4). Although we used sex- and age-normalized T-scores according to the NEO-PI-R manual, we observed some remaining associations between the NEO personality traits and both sex and age. Detailed association results between covariates and our outcome variables are shown in supplementary Table S3.

### Big Five personality traits and CNS arousal

Spearman correlations between the five NEO personality dimensions (sex and age-normalized T-scores) and the three EEG-vigilance variables are shown in Table 2.

**Table 2.** Spearman correlations between NEO personality dimensions (T-Scores) and EEG-vigilance variables

| $N = 468$ | Mean vigilance | | | | Stability score | | | | Slope index | | | |
| --- | --- | --- | --- | --- | --- | --- | --- | --- | --- | --- | --- | --- |
| | rho | $p$ | FDR | BF<sub>10</sub> | rho | $p$ | FDR | BF<sub>10</sub> | rho | $p$ | FDR | BF<sub>10</sub> |
| Neuroticism | -.063 | .170 | .365 | 0.27 | -.030 | .515 | .766 | 0.13 | -.002 | .972 | .972 | 0.11 |
| Extraversion | -.104 | .025* | .082 | 1.29 | -.096 | .038* | .095 | 0.91 | -.137 | .003* | .023** | 8.35 |
| Openness | -.102 | .027* | .082 | 1.20 | -.121 | .009* | .044** | 3.23 | -.173 | 2E-4* | .002** | 121.33 |
| Agreeableness | -.027 | .561 | .766 | 0.13 | -.012 | .791 | .913 | 0.11 | -.032 | .496 | .766 | 0.14 |
| Conscientiousness | -.023 | .624 | .780 | 0.12 | -.005 | .913 | .972 | 0.11 | -.039 | .399 | .748 | 0.15 |

FDR: False Discovery Rate according to Benjamini and Hochberg; BF<sub>10</sub> Bayes factor showing the likelihood ratio between the alternate and null hypothesis (1/3 beta prior width).
\* $p < .05$ (two-sided nominal significance)
\*\* FDR < .05 (p-value corrected for all tested associations using FDR method)

Analyses revealed six associations with nominal significance (*p* < .05). Of these, three remained significant after multiple testing correction (FDR < .05). We observed EEG-vigilance to be inversely associated with the degree of extraversion (slope index: r_S_ = -.137, p = .003, FDR = .023, BF_10_ = 8.35) and openness to experience (stability score: r_S_ = -.121, p = .009, FDR = .044, BF_10_ = 3.23; slope index: r_S_ = -.173, p = 2E-4, FDR = .002, BF_10_ = 121.33). Subjects who reported higher levels of extraversion and openness to experience, respectively, exhibited lower EEG-vigilance. For illustrative purposes, the time-courses of EEG-vigilance stratified by groups scoring low vs. high on the respective Big Five dimension (lower vs. upper quartile of the ascending distribution) are shown in Figure 1.

**Fig. 1.**
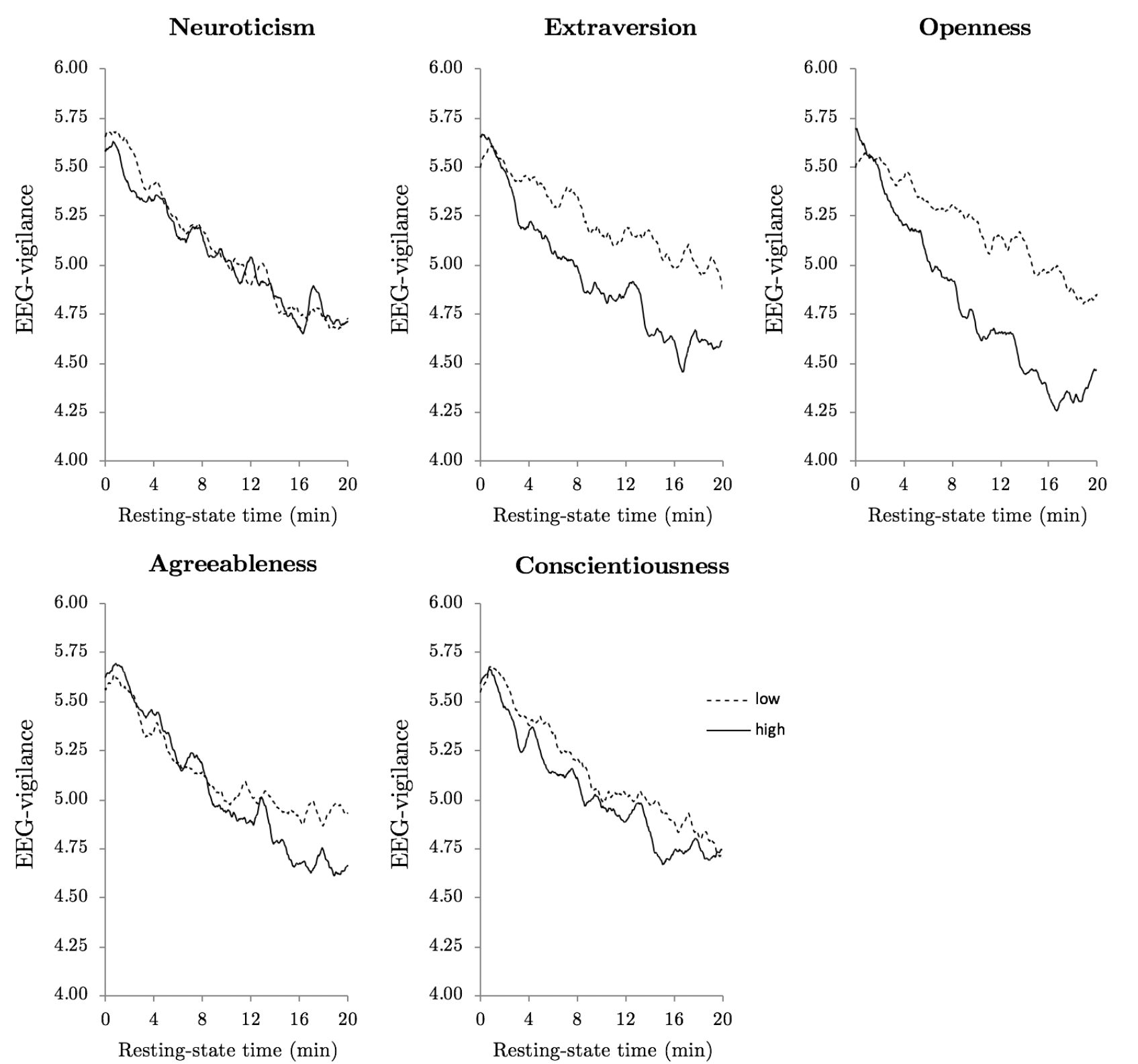
Time-courses of EEG-vigilance during the 20-minute eyes-closed resting-state condition stratified by groups scoring low vs. high on the respective Big Five scale (i.e. subjects with scores in the lower vs. upper quartile of the ascending distribution). Time-courses reflect simple moving averages (SMA), i.e., every data point represents an averaged 61-second interval of EEG-vigilance (data point in time ± 30 seconds). Statistical analyses revealed significant correlations between EEG-vigilance and both extraversion and openness to experience.

We repeated our analysis by additionally adjusting correlations by sex, age, and daytime of EEG assessment (suppl. Table S4). This resulted in three associations reaching nominal significance. Of these, one remained significant after multiple testing correction (in this case the FDR corrected p-value is equivalent to the Bonferroni-corrected p-value): Subjects who reported higher levels of openness to experience exhibited lower EEG-vigilance (slope index: r**S** = -.152, p = .001, FDR = .015, BF_10_ = 23.40). In order to examine whether the distribution of observed p-values differs from a random p-value distribution, we generated a permutation-based qq-plot that takes into account the dependencies between association tests (Fig. 2A).

**Fig. 2.**
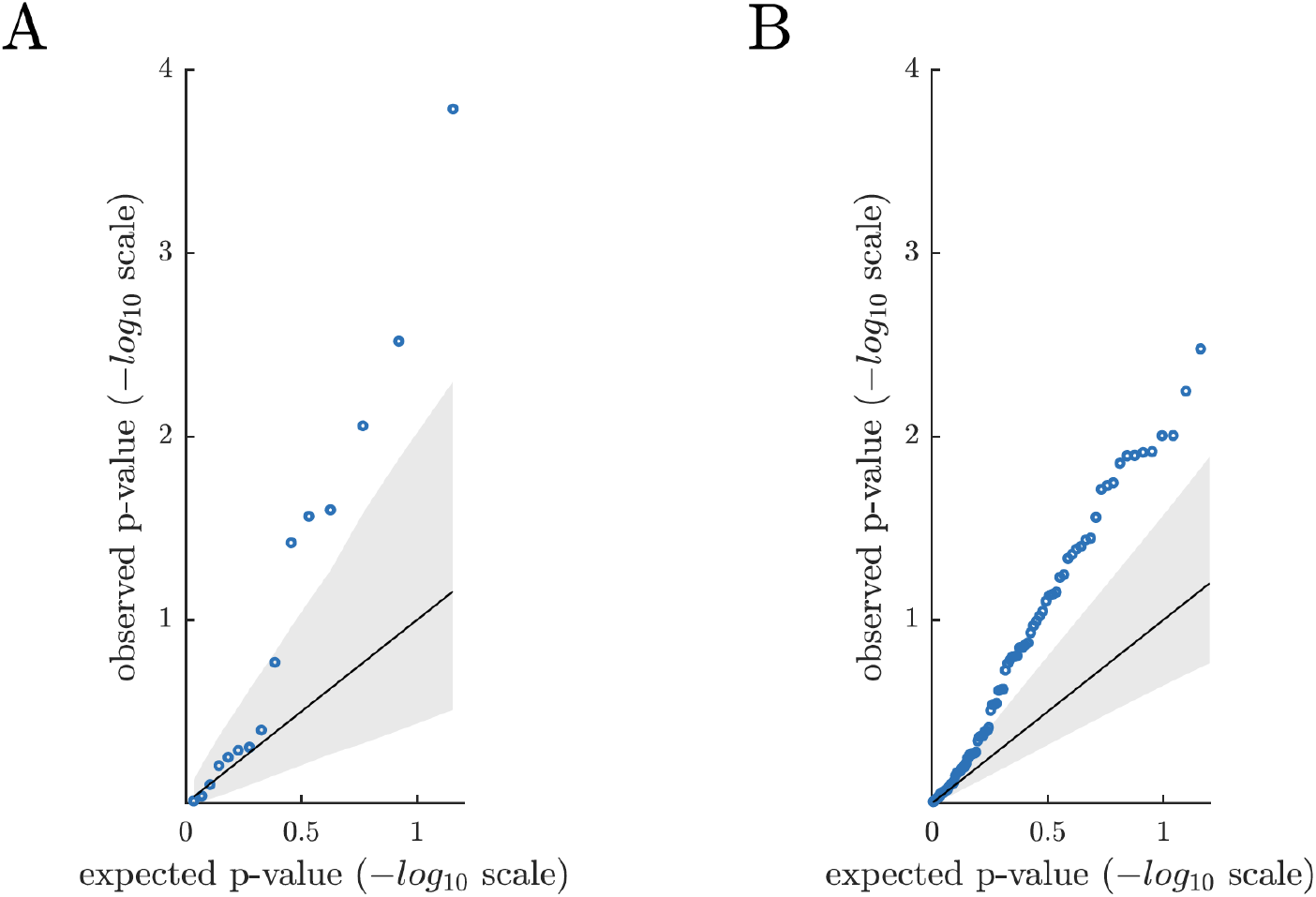
Permutation-based qq-plot showing the observed p-values from the association analyses (blue circles) plotted against the expected p-values under the null hypothesis. The solid diagonal line represents the mean expected p-values. The lower and upper bound of the grey area represent the 5^th^ and 95^th^ percentile (-log10 scale) of the expected p-values. **A** Results based on the sex- and age-normalized T-scores of the NEO personality traits. **B** Results based on the sex- and age-normalized T-scores of the NEO personality facets. In total, qq-plots suggest that association analyses revealed stronger evidence than expected by chance.

The qq-plot shows that the six strongest observed p-values exceed the 95^th^ percentile (-log10 scale) of the computed expected p-value distribution. In detail, only 0.7% of the one million sets of expected p-values contained at least six p-values below 0.038 (that is the 6th lowest observed p-value), and 0.2% of the one million sets of expected p-values contained at least one p-value below 2E-4 (that is the lowest observed p-value). Overall, the plot indicates that the distribution of observed p-values differs from a random p-value distribution under the null hypothesis. When additionally adjusting association results by sex, age, and daytime of EEG assessment, only the strongest observed association exceeded the 95^th^ percentile of the expected p-value distribution (suppl. Figure S3A). In this regard, only 1.1% of the one million sets of expected p-values contained one p-value lower than 0.001 (that is the lowest observed p-value).

### NEO personality facets and CNS arousal

To further elaborate the nature of the underlying associations, we carried out exploratory Spearman correlations between the 30 NEO facets (sex and age-normalized T-scores) and each of the 3 EEG-vigilance variables. Detailed association results are shown in supplementary Table S5. In total, 24 out of 90 correlations reached the level of nominal significance. The strongest association was observed for neuroticism facet ‘impulsiveness’ (mean vigilance: r_S_ = -.150, p = .001, BF_10_ = 19.88). No other neuroticism facet reached nominal significance. Regarding extraversion, we found nominally significant results for the facets ‘warmth’ (slope index: r_S_ = -.119, p = .010, BF_10_ = 2.90), ‘assertiveness’ (slope index: r_S_ = -.109, p = .018, BF_10_ = 1.73), ‘activity’ (slope index: r_S_ = -.114, p = .014, BF_10_ = 2.14), and ‘positive emotions’ (slope index: r_S_ = -.116, p = .012, BF_10_ = 2.43). Regarding openness to experience, we found significant associations for the facets ‘fantasy’ (slope index: r_S_ = -.094, p = .041, BF_10_ = 0.85), ‘aesthetics’ (slope index: r_S_ = -.137, p = .003, BF_10_ = 8.60), ‘feelings’ (slope index: r_S_ = -.137, p = .003, BF_10_ = 8.27), ‘actions’ (slope index: r_S_ = -.109, p = .018, BF_10_ = 1.68), and ‘ideas’ (slope index: r_S_ = -.128, p = .006, BF_10_ = 4.76). We also observed nominally significant results for agreeableness facet ‘tender-mindedness’ (slope index: r_S_ = -.145, p = .002, BF_10_ = 14.40) and conscientiousness facet ‘achievement striving’ (slope index: r_S_ = -.135, p = .003, BF_10_ = 7.65).

By analogy to the NEO personality dimension analyses, we generated a permutation-based qq-plot to examine whether the distribution of observed p-values of the NEO personality facets differs from a random p-value distribution (Fig. 2B). Again, the qq-plot indicates that association analyses revealed stronger evidence than expected by chance. Notably, consistent with the Big Five results, we observed an attenuation of effect sizes when additionally adjusting sex- and age-normalized T-Score correlations by sex, age and daytime of EEG-assessment (suppl. Table S6). The distribution of observed vs. expected p-values after adjusting T-Scores is shown in supplementary Figure S3B, with a large proportion still exceeding the 95^th^ percentile of expected p-values.

## Discussion

In this study, we investigated the association between the Big Five personality traits and CNS arousal in the resting-state by making use of the EEG- and EOG-based analysis tool VIGALL. Our primary analysis suggests that, after multiple testing correction, CNS arousal is negatively associated with the degree of extraversion and openness to experience: Subjects who reported higher levels of extraversion and openness to experience, respectively, showed steeper declines of EEG-vigilance. In addition, when considering all tested associations between the Big Five personality traits and CNS arousal, we observed overall stronger effects than expected by chance. This finding was supported by association results of the thirty NEO personality facets. Facet analyses also revealed that the observed associations of the higher-order Big Five traits and CNS arousal were not driven by a single facet with a distinct, strong effect but rather appeared to arise from a distributed pattern of associations across several facets. Notably, for the majority of nominally significant associations (p < 0.05), Bayesian analysis revealed only anecdotal evidence for the alternate hypothesis (BF_10_ ranging between 1 and 3). In addition, when taking into account potential confounders, we observed a general attenuation of effect sizes, with several associations dropping below the nominal and FDR-corrected level of significance. Further, we did not obtain evidence for an association of CNS arousal and neuroticism, a personality trait that we regard as highly plausible candidate for arousal alterations. Overall, across frequentist and Bayesian analyses and irrespective of accounting for potential confounders or not, we obtained the strongest and most compelling evidence for a link between openness to experience and CNS arousal.

To our knowledge, this investigation is the largest EEG study so far addressing the link between CNS arousal and the Big Five personality traits. In keeping with this, the statistical power to detect associations in this study was substantially higher when compared to the vast majority of previous investigations, which usually featured a sample size fewer than 100 subjects. Given the present study design and analysis procedure, the achieved statistical power enables to conclude that neuroticism is unlikely to account for more than 4% (r_S_ ≥ 0.2) of the variance in CNS arousal, since the probability (1-β) of identifying such an effect at p < 0.05 exceeded 99.98% (see suppl. Fig. S1 for a power plot). Similarly, extraversion surpassed the FDR-corrected but not the Bonferroni-adjusted level of significance, with the latter being reached with a probability above 92% given a true effect size of r ≥ 0.20. Thus, if extraversion and neuroticism are truly associated with CNS arousal, correlations are certainly below r = 0.20. This is well in agreement with a study of 708 meta-analytically derived correlations in the field of personality and individual differences research, suggesting a median reported effect size of r = 0.19 (Gignac & Szodorai, 2016). Notably, when considering preregistered studies only, the median effect size has been reported to be even lower (r = 0.12; Schäfer & Schwarz, 2019). Accordingly, to elucidate the biological basis of individual differences in human personality, we believe that there is an urgent need for large (collaborative) studies with at least several hundreds and preferably thousands of subjects.

The present study adds empirical results to the ongoing debate of whether extraverted individuals exhibit lower habitual levels of CNS arousal. Consistent with the theoretical assumptions, our primary analyses provided supportive evidence for a negative correlation between extraversion and arousal. Nevertheless, there remain some reservations that we would like to outline. First, although we used sex- and age-normalized T-scores, we still observed associations between the NEO personality scores and both sex and age (suppl. Table S3). After considering sex and age as additional covariates, the observed associations between extraversion and CNS arousal did not remain significant after multiple testing correction. Hence, the present results do not provide stringent support for extraversion to share unique variance with CNS arousal beyond the effects of sex and age. Second, Bayesian analyses provided only anecdotal to moderate evidence for the proposed link. This may partly be explained by the selected priors. Here, we used a symmetrically scaled 1/3 beta prior, which is the default setting of R package ‘BayesFactor’ (Morey & Rouder, 2018). This prior corresponds to the expectation that with an 80% probability the true effect falls in between r = −0.5 and r = 0.5. However, in light of the reported effect sizes in the field of individual differences research (Gignac & Szodorai, 2016; Schäfer & Schwarz, 2019), this prior width might still be considered too wide, and resulting Bayes factors may thus show some bias towards favoring the null hypothesis. Notably, the software package JASP, which is widely used in the social and behavioral sciences, implicates an even more naive uniform prior as default setting (JASP Team, 2020). Hence, given the rising popularity of Bayesian analyses in the life sciences, we feel that the selection of adequate prior widths may be one crucial topic of the future scientific debate. Taken together, we here find some evidence supporting Eysenck’s postulations concerning the link between extraversion and CNS arousal, but the observed effect strength suggests that even larger sample sizes are required to establish reliable associations that withstand a rigorous control for potential confounders. An elaborated a priori knowledge of the expected effect sizes may further increase the study power.

Although we did not obtain evidence for a link between CNS arousal and neuroticism, exploratory analyses revealed indications for a negative association with neuroticism facet ‘impulsiveness’. This result was the most compelling among all facet associations and remained significant after multiple-testing correction. Interestingly, impulsiveness showed relatively low correlations with the other neuroticism facets and, in contrast to them, correlated positively with facets of extraversion and openness to experience (suppl. Fig. S2). In this light, impulsiveness may be considered as rather atypical facet of the higher-order trait neuroticism. Notably, the observation of low habitual arousal levels in individuals exhibiting impulsive behavior is well in line with previously proposed concepts (Eysenck, 1967; Hegerl & Hensch, 2014; Zuckerman, 1979).

In comparison to neuroticism facet ‘impulsiveness’, our analyses did not reveal indications for a link between neuroticism facet ‘depression’ and CNS arousal. By applying the Vigilance Algorithm Leipzig, several previous studies provided supportive evidence for an association between clinical depression and enhanced CNS arousal in the resting state (Hegerl et al., 2012; Sander et al., 2018; Schmidt et al., 2016, 2017; Ulke et al., 2017; Ulke, Tenke, et al., 2019; Ulke, Wittekind, et al., 2019; Wittekind et al., 2016). Although the present study included subjects without a current depression diagnosis, it can be assumed that alterations in CNS arousal occur in both the normal and pathological range of human behavior. This is consistent with the view that personality traits and psychopathology are no distinct entities, but may rather manifest along a common spectrum of functioning (Widiger, 2011). This argumentation also ties in with the postulations of the Research Domain Criteria Project (RDoC), according to which mental diseases can be considered to fall along multiple continuous trait dimensions, with traits ranging from normal to the extreme (Cuthbert & Insel, 2013). Intriguingly, the RDoC project considers ‘arousal and regulatory systems’ as one out of five fundamental domains to describe and classify psychiatric disorders. In the present study, the lack of evidence for an association between CNS arousal and ‘depression’ may be explained by lower effect sizes among healthy subjects relative to the study of healthy control vs. in-patient samples.

Across all analyses, we obtained the strongest evidence for an association between the Big Five personality trait openness to experience and CNS arousal. Bayes factors indicated ‘extreme evidence’ (BF_10_ > 100) and ‘strong evidence’ (BF_10_ ranging from 10 to 30) for this link, respectively, depending on whether we considered zero-order correlations or whether sex, age, and daytime of EEG assessment served as covariates. Interestingly, previous genetic correlation analyses suggest a positive association between openness to experience and major depression as well as bipolar disorder (Lo et al., 2017). These mental diseases have both been argued to possess arousal-related pathophysiologies (Hegerl & Hensch, 2014). However, since manic and depressive-like behavior have been postulated to be linked to habitually low vs. high arousal levels, it remains difficult to deduce the sign of the potential association between openness to experience and CNS arousal. Importantly, previous investigations have shown that openness to experience positively correlates with sensation seeking (Aluja, García, & García, 2003). Further, openness to experience has been reported to positively correlate with extraversion, which is also shown in the present dataset (suppl. Table S1). On this account, we regard the present findings of lower arousal levels in subjects scoring high on openness to experience as consistent with the concepts of Eysenck (1967) and Zuckerman (1979).

Our study poses some limitations that need to be addressed. First, our participants were, on average, 58 years old, and reported association strengths may not generalize across other age groups. In particular, we previously observed a general tendency towards stronger effect sizes for arousal associations among the younger age groups (Jawinski et al., 2017). This might be explained be the higher EEG total power in younger adults and a possibly related higher accuracy of EEG-vigilance classifications. Further, we here addressed the relationship between individual differences in personality and habitual levels of CNS arousal by means of an EEG resting-state paradigm. However, one major emphasis of Eysenck’s theory is put on the differential performance of extraverts and introverts as a function of arousal-enhancing situational factors (Brocke & Battmann, 1992). Thus, an interesting future direction might be the use of VIGALL in experimental studies with behavioral performance outcomes (e.g., as done previously by Huang et al., 2017) while taking into account the ‘hedonic tone of an individual’, i.e., the preferred level of excitation. Lastly, it should be noted that the present study sought to answer the question of whether CNS arousal but not the EEG, in general, is predictive for basic personality traits. Stronger associations may be derived by the application of machine learning models trained on the EEG to directly predict human personality (for an example see Li et al., 2019).

## Conclusion

To the best of our knowledge, this study is the largest EEG study so far addressing the relationship between the Big Five personality traits and habitual levels of CNS arousal. Concerning Eysenck’s Arousal Activation Theory, our results provide some support for extraversion and no support for neuroticism to be linked to CNS arousal. Intriguingly, Bayesian and frequentist analyses revealed convincing evidence for a link between openness to experience and lower levels of CNS arousal. In addition, among the lower-order personality traits, we obtained evidence for neuroticism facet ‘impulsivity’ and reduced CNS arousal. We regard these findings as well in line with the postulations of Eysenck and Zuckerman and consistent with the assumptions of the arousal regulation model. In total, the present study results agree with meta-analytically derived effect sizes in the field of personality and individual differences research, highlighting the need for large (collaborative) studies with at least several hundreds of subjects.

## Acknowledgements

This publication is supported by LIFE – Leipzig Research Center for Civilization Diseases, Universität Leipzig. LIFE is funded by means of the European Union, by the European Regional Development Fund (ERDF) and by means of the Free State of Saxony within the framework of the excellence initiative. This project was funded by means of the European Social Fund and the Free State of Saxony. This study was supported within the framework of the cooperation between the German Depression Foundation and the ‘Deutsche Bahn Stiftung gGmbH’.

## Supplementary Material

Supplementary information is available at bioRxiv online.

