## Supplemental Material for "The Big Five personality traits and CNS arousal in the resting state"

bioRxiv

**Power analysis and intercorrelations**

| | |
|---|---|
| Supplementary Fig. S1 | Power analysis results |
| Supplementary Table S1 | NEO personality dimensions (T-scores) - Cronbach's Alpha and intercorrelations |
| Supplementary Fig. S2 | NEO personality facets (T-scores) - Cronbach's Alpha and intercorrelations |
| Supplementary Table S2 | Intercorrelations between EEG-vigilance variables |
| Supplementary Table S3 | Correlations of the covariates sex, age, and daytime of EEG assessment with NEO personality traits (T-scores) and EEG-vigilance |

**Associations between NEO personality traits and EEG-vigilance**

| | |
|---|---|
| Supplementary Fig. S3 | Permutation-based qq-plot of observed vs. expected p values for NEO personality traits and facets (T-Scores) after adjusting for covariates |
| Supplementary Table S4 | Partial Spearman correlations between NEO personality dimensions (T-Scores) and EEG-vigilance variables |
| Supplementary Table S5 | Spearman correlations between NEO personality facets (T-Scores) and EEG-vigilance variables |
| Supplementary Table S6 | Partial Spearman correlations between NEO personality facets (T-Scores) and EEG-vigilance variables |

### Power analysis and psychometric properties

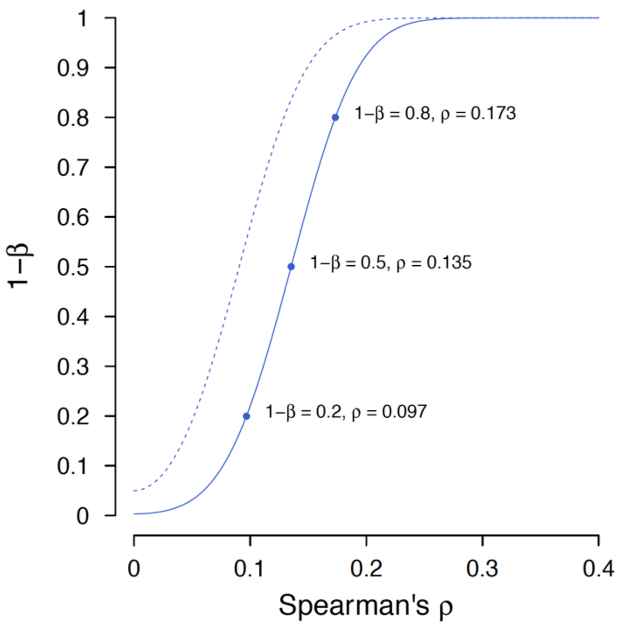


**Fig. S1** Power analysis results showing the probability (1-β) of associations to surpass the threshold of significance given true effect sizes ranging between ρ = 0.0 and ρ = 0.4 (with $N$ = 468). The dotted curve shows the probability to reach nominal significance (α = 0.05, two-sided). The solid curve shows the probability to reach the Bonferroni-corrected level of significance (α = 0.05/15, two-sided). Power analysis was conducted using R package pwr v1.2-2 (Champely et al., 2018).

**Table S1** NEO personality traits (T-Scores) – Cronbach's α and intercorrelations

| $N$ = 468 | N | | E | | O | | A | | C | |
|---|---|---|---|---|---|---|---|---|---|---|
| | rho | $p$ | rho | $p$ | rho | $p$ | rho | $p$ | rho | $p$ |
| N Neuroticism | .906 | - | | | | | | | | |
| E Extraversion | -.337 | 7E-14* | .899 | - | | | | | | |
| O Openness | -.249 | 5E-8* | .477 | 5E-28* | .868 | - | | | | |
| A Agreeableness | -.190 | 4E-5* | -.069 | .136 | .071 | .127 | .836 | - | | |
| C Conscientiousness | -.440 | 1E-23* | .333 | 1E-13* | .121 | .009* | .200 | 1E-5* | .881 | - |

Results show Spearman correlations. Values of the main diagonal reflect the internal consistency (Cronbach's α).

* $p < .05$ (two-sided nominal significance)

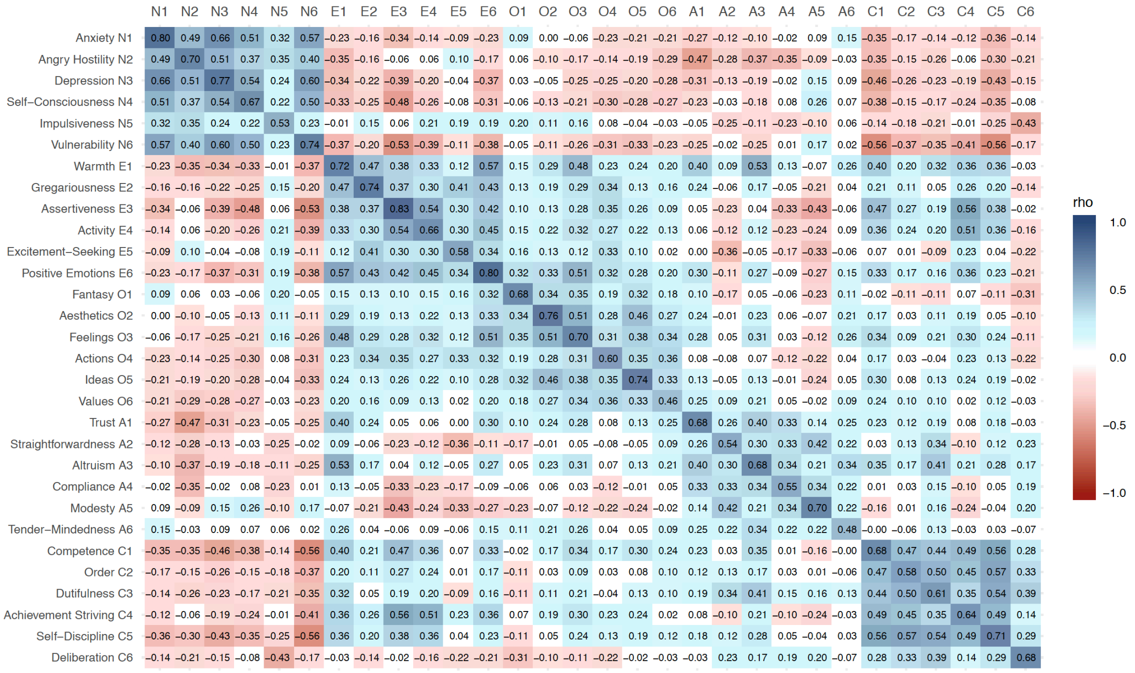


**Fig. S2** NEO personality facets - Cronbach's α and intercorrelations. Only cells containing correlations with nominal significance (rho > 0.0906, $p < 0.05$) have been assigned with colors of the blue and red color palette. Values of the main diagonal reflect the internal consistency as estimated using Cronbach's α.

**Table S2** Intercorrelations between EEG-vigilance correlations

| $N = 468$ | Mean vigilance | | Stability score | | Slope index | |
|---|---|---|---|---|---|---|
| | rho | $p$ | rho | $p$ | rho | $p$ |
| Mean vigilance | 1.000 | - | | | | |
| Stability score | .821 | 1E-115* | 1.000 | - | | |
| Slope index | .820 | 5E-115* | .881 | 9E-154 | 1.000 | - |

Results show Spearman correlations.

* $p < .05$ (two-sided nominal significance)

**Table S3** Correlations of sex, age, and daytime of EEG assessment with NEO personality traits (T-scores) and EEG-vigilance variables

| $N = 468$ | Sex | | Age | | Daytime | |
|---|---|---|---|---|---|---|
| | rho | $p$ | rho | $p$ | rho | $p$ |
| **EEG-vigilance** | | | | | | |
| Mean vigilance | .021 | .648 | .168 | 3E-4* | -.155 | 8E-4* |
| Stability score | .030 | .523 | .178 | 1E-4* | -.197 | 2E-5* |
| Slope index | .084 | .069 | .191 | 3E-5* | -.167 | 3E-4* |
| **NEO personality dimensions** | | | | | | |
| Neuroticism | .064 | .164 | -.028 | .539 | .017 | .722 |
| Extraversion | -.161 | 5E-4* | -.074 | .111 | .068 | .143 |
| Openness | -.076 | .102 | -.084 | .070 | .012 | .792 |
| Agreeableness | .009 | .840 | -.074 | .109 | -.070 | .131 |
| Conscientiousness | -.164 | 4E-4* | .080 | .082 | -.020 | .662 |
| **NEO personality facets** | | | | | | |
| Neuroticism | | | | | | |
| N1 Anxiety | .028 | .539 | -.024 | .606 | .010 | .825 |
| N2 Angry Hostility | .076 | .099 | .012 | .803 | .040 | .383 |
| N3 Depression | .047 | .314 | .025 | .597 | .018 | .692 |
| N4 Self-Consciousness | .085 | .067 | -.060 | .199 | -.048 | .298 |
| N5 Impulsiveness | -.050 | .279 | .012 | .801 | .048 | .296 |
| N6 Vulnerability | .114 | .013* | -.066 | .154 | -.039 | .400 |
| Extraversion | | | | | | |
| E1 Warmth | -.115 | .012* | -.093 | .044* | .087 | .059 |
| E2 Gregariousness | -.032 | .484 | -.140 | .002* | .009 | .842 |
| E3 Assertiveness | -.078 | .092 | .024 | .598 | .087 | .060 |
| E4 Activity | -.142 | .002* | -.014 | .765 | .067 | .148 |
| E5 Excitement-Seeking | -.095 | .040* | -.050 | .279 | .020 | .659 |
| E6 Positive Emotions | -.233 | 4E-7* | -.061 | .190 | .042 | .366 |
| Openness | | | | | | |
| O1 Fantasy | -.060 | .192 | -.065 | .159 | .005 | .908 |
| O2 Aesthetics | -.049 | .289 | -.055 | .237 | .038 | .411 |
| O3 Feelings | -.108 | .020* | -.101 | .029* | .041 | .375 |
| O4 Actions | -.039 | .398 | -.049 | .288 | .045 | .333 |
| O5 Ideas | -.018 | .702 | -.010 | .833 | -.030 | .521 |
| O6 Values | -.035 | .452 | .000 | .998 | -.069 | .137 |
| Agreeableness | | | | | | |
| A1 Trust | -.035 | .452 | -.074 | .112 | -.008 | .861 |
| A2 Straightforwardness | .038 | .413 | -.015 | .754 | -.036 | .431 |
| A3 Altruism | -.103 | .026* | -.069 | .137 | -.037 | .423 |
| A4 Compliance | .123 | .008* | -.024 | .604 | -.055 | .239 |
| A5 Modesty | .006 | .897 | .031 | .505 | -.073 | .115 |
| A6 Tender-Mindedness | -.002 | .969 | -.165 | 4E-4* | -.042 | .359 |
| Conscientiousness | | | | | | |
| C1 Competence | -.129 | .005* | .040 | .384 | .026 | .581 |
| C2 Order | -.119 | .010* | .087 | .061 | -.048 | .299 |
| C3 Dutifulness | -.184 | 6E-5* | -.013 | .775 | -.021 | .653 |
| C4 Achievement Striving | -.129 | .005* | .062 | .180 | .052 | .264 |
| C5 Self-Discipline | -.093 | .045* | .039 | .400 | -.050 | .285 |
| C6 Deliberation | -.050 | .278 | .103 | .026* | -.052 | .261 |

Results show Spearman correlations. Sex was coded as male = 1 and female = 2.

* $p < .05$ (two-sided nominal significance)

#### Associations between NEO personality traits and EEG-vigilance

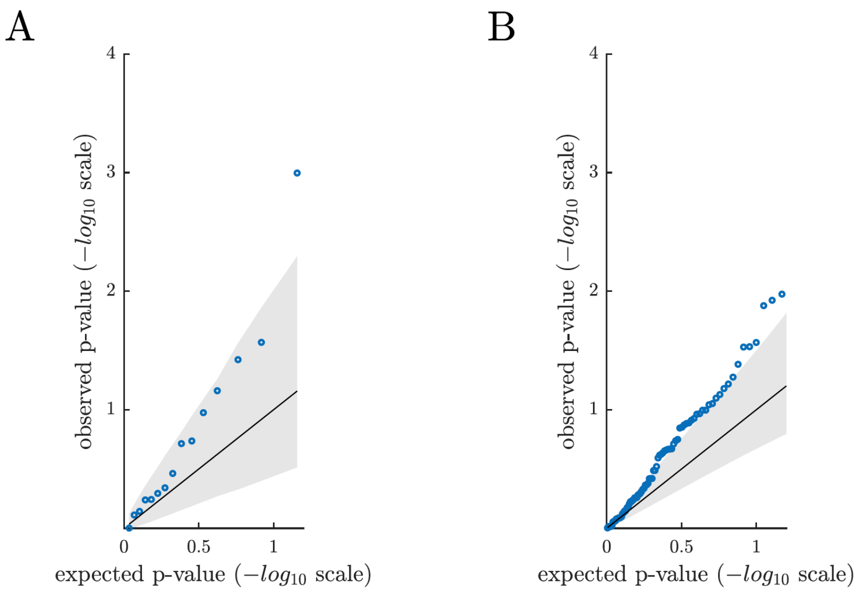


**Fig. S3** Permutation-based qq-plot showing the observed p values from the association analyses (blue circles) plotted against the expected p values under the null hypothesis. The solid diagonal line represents the mean expected p values. The lower and upper bound of the grey area represent the 5$^{th}$ and 95$^{th}$ percentile (-log10 scale) of the expected p values. **A** Association results of the higher-order NEO personality traits (T-scores) additionally adjusted by sex, age, and time of EEG assessment. **B** Association results based on the sex- and age-normalized T scores of the NEO personality facets.

**Table S4** Partial Spearman correlations between NEO personality dimensions (T-Scores) and EEG-vigilance variables

| $N = 468$ | Mean vigilance | | | | Stability score | | | | Slope index | | | |
|---|---|---|---|---|---|---|---|---|---|---|---|---|
| | rho | $p$ | FDR | $BF_{10}$ | rho | $p$ | FDR | $BF_{10}$ | rho | $p$ | FDR | $BF_{10}$ |
| Neuroticism | -.060 | .193 | .414 | 0.25 | -.026 | .577 | .721 | 0.13 | .000 | .997 | .997 | 0.11 |
| Extraversion | -.075 | .106 | .318 | 0.40 | -.062 | .184 | .414 | 0.26 | -.096 | .038* | .189 | 0.92 |
| Openness | -.084 | .069 | .260 | 0.56 | -.103 | .027* | .189 | 1.23 | -.152 | .001* | .015** | 23.40 |
| Agreeableness | -.026 | .572 | .721 | 0.13 | -.013 | .773 | .828 | 0.11 | -.031 | .508 | .721 | 0.13 |
| Conscientiousness | -.035 | .456 | .721 | 0.14 | -.017 | .721 | .828 | 0.12 | -.044 | .345 | .647 | 0.17 |

Effects of sex, age, and daytime of EEG assessment were partialled out. FDR: False Discovery Rate according to Benjamini and Hochberg; $BF_{10}$ Bayes factor showing the likelihood ratio between the alternate and null hypothesis (1/3 beta prior width).

* $p < .05$ (two-sided nominal significance)
** FDR $< .05$ ($p$ value corrected for all tested associations using FDR method)

**Table S5** Spearman correlations between NEO personality facets (T-Scores) and EEG-vigilance variables

| $N = 468$ | Mean vigilance | | | | Stability score | | | | Slope index | | | |
|---|---|---|---|---|---|---|---|---|---|---|---|---|
| | rho | $p$ | FDR | $BF_{10}$ | rho | $p$ | FDR | $BF_{10}$ | rho | $p$ | FDR | $BF_{10}$ |
| Neuroticism | | | | | | | | | | | | |
| N1 Anxiety | -.049 | .290 | .512 | 0.19 | -.029 | .533 | .768 | 0.13 | -.020 | .667 | .854 | 0.12 |
| N2 Angry Hostility | -.054 | .243 | .455 | 0.21 | -.028 | .544 | .768 | 0.13 | .004 | .928 | .971 | 0.11 |
| N3 Depression | -.007 | .885 | .949 | 0.11 | .007 | .877 | .949 | 0.11 | .022 | .638 | .844 | 0.12 |
| N4 Self-Consciousness | -.068 | .141 | .327 | 0.32 | -.021 | .648 | .845 | 0.12 | .008 | .868 | .949 | 0.11 |
| N5 Impulsiveness | -.150 | .001* | .050** | 19.88 | -.076 | .101 | .276 | 0.41 | -.087 | .059 | .203 | 0.63 |
| N6 Vulnerability | .017 | .707 | .871 | 0.12 | .009 | .850 | .949 | 0.11 | .063 | .172 | .353 | 0.27 |
| Extraversion | | | | | | | | | | | | |
| E1 Warmth | -.116 | .012* | .088 | 2.41 | -.097 | .036* | .165 | 0.95 | -.119 | .010* | .088 | 2.90 |
| E2 Gregariousness | -.019 | .677 | .854 | 0.12 | .007 | .874 | .949 | 0.11 | -.005 | .917 | .971 | 0.11 |
| E3 Assertiveness | -.081 | .079 | .237 | 0.50 | -.083 | .073 | .229 | 0.53 | -.109 | .018* | .103 | 1.73 |
| E4 Activity | -.102 | .028* | .138 | 1.19 | -.079 | .090 | .260 | 0.45 | -.114 | .014* | .090 | 2.14 |
| E5 Excitement-Seeking | -.024 | .607 | .828 | 0.12 | -.065 | .158 | .340 | 0.29 | -.037 | .430 | .675 | 0.15 |
| E6 Positive Emotions | -.083 | .074 | .229 | 0.53 | -.065 | .159 | .340 | 0.29 | -.116 | .012* | .088 | 2.43 |
| Openness | | | | | | | | | | | | |
| O1 Fantasy | -.084 | .071 | .229 | 0.54 | -.077 | .095 | .268 | 0.43 | -.094 | .041* | .168 | 0.85 |
| O2 Aesthetics | -.092 | .046* | .173 | 0.77 | -.097 | .037* | .165 | 0.93 | -.137 | .003* | .050** | 8.60 |
| O3 Feelings | -.108 | .019* | .103 | 1.61 | -.093 | .044* | .172 | 0.80 | -.137 | .003* | .050** | 8.27 |
| O4 Actions | -.034 | .457 | .697 | 0.14 | -.075 | .107 | .284 | 0.39 | -.109 | .019* | .103 | 1.68 |
| O5 Ideas | -.068 | .142 | .327 | 0.31 | -.095 | .040* | .168 | 0.87 | -.128 | .006* | .073 | 4.76 |
| O6 Values | .039 | .406 | .664 | 0.15 | .014 | .759 | .923 | 0.11 | -.002 | .971 | .980 | 0.11 |
| Agreeableness | | | | | | | | | | | | |
| A1 Trust | -.069 | .133 | .327 | 0.33 | -.029 | .536 | .768 | 0.13 | -.069 | .138 | .327 | 0.32 |
| A2 Straightforwardness | -.002 | .957 | .979 | 0.11 | -.013 | .783 | .931 | 0.11 | .009 | .841 | .949 | 0.11 |
| A3 Altruism | -.001 | .980 | .980 | 0.11 | -.037 | .428 | .675 | 0.15 | -.055 | .239 | .455 | 0.21 |
| A4 Compliance | .050 | .285 | .512 | 0.19 | .040 | .387 | .656 | 0.16 | .029 | .526 | .768 | 0.13 |
| A5 Modesty | .054 | .240 | .455 | 0.21 | .088 | .057 | .203 | 0.65 | .072 | .118 | .302 | 0.36 |
| A6 Tender-Mindedness | -.146 | .002* | .050** | 15.16 | -.119 | .010* | .088 | 2.90 | -.145 | .002* | .050** | 14.40 |
| Conscientiousness | | | | | | | | | | | | |
| C1 Competence | -.047 | .312 | .540 | 0.18 | -.027 | .566 | .783 | 0.13 | -.066 | .157 | .340 | 0.29 |
| C2 Order | .012 | .801 | .936 | 0.11 | .022 | .635 | .844 | 0.12 | -.013 | .787 | .931 | 0.11 |
| C3 Dutifulness | -.036 | .435 | .675 | 0.15 | -.011 | .819 | .945 | 0.11 | -.039 | .403 | .664 | 0.15 |
| C4 Achievement Striving | -.115 | .013* | .088 | 2.33 | -.115 | .013* | .088 | 2.32 | -.135 | .003* | .050** | 7.65 |
| C5 Self-Discipline | -.019 | .684 | .854 | 0.12 | .003 | .943 | .975 | 0.11 | -.028 | .546 | .768 | 0.13 |
| C6 Deliberation | .064 | .165 | .345 | 0.28 | .061 | .188 | .376 | 0.25 | .049 | .289 | .512 | 0.19 |

FDR: False Discovery Rate according to Benjamini and Hochberg; $BF_{10}$: Bayes factor showing the likelihood ratio between the alternate and null hypothesis (1/3 beta prior width)

* $p < .05$ (two-sided nominal significance)
** FDR $< .05$ ($p$ value corrected for all tested associations using FDR method)

**Table S6** Partial Spearman correlations between NEO personality facets (T-Scores) and EEG-vigilance variables

| $N = 468$ | Mean vigilance | | | | Stability score | | | | Slope index | | | |
|---|---|---|---|---|---|---|---|---|---|---|---|---|
| | rho | $p$ | FDR | $BF_{10}$ | rho | $p$ | FDR | $BF_{10}$ | rho | $p$ | FDR | $BF_{10}$ |
| Neuroticism | | | | | | | | | | | | |
| N1 Anxiety | -.046 | .326 | .652 | 0.17 | -.025 | .596 | .840 | 0.12 | -.017 | .716 | .923 | 0.12 |
| N2 Angry Hostility | -.055 | .238 | .531 | 0.22 | -.028 | .554 | .830 | 0.13 | .000 | .994 | .994 | 0.11 |
| N3 Depression | -.011 | .820 | .951 | 0.11 | .004 | .937 | .984 | 0.11 | .015 | .743 | .941 | 0.11 |
| N4 Self-Consciousness | -.072 | .123 | .429 | 0.35 | -.026 | .582 | .840 | 0.13 | .002 | .962 | .984 | 0.11 |
| N5 Impulsiveness | -.146 | .002* | .126 | 15.46 | -.068 | .143 | .429 | 0.31 | -.079 | .089 | .427 | 0.46 |
| N6 Vulnerability | .018 | .693 | .917 | 0.12 | .007 | .874 | .959 | 0.11 | .060 | .194 | .527 | 0.25 |
| Extraversion | | | | | | | | | | | | |
| E1 Warmth | -.083 | .075 | .419 | 0.53 | -.057 | .219 | .527 | 0.23 | -.075 | .108 | .427 | 0.39 |
| E2 Gregariousness | .010 | .835 | .951 | 0.11 | .041 | .381 | .715 | 0.16 | .032 | .488 | .813 | 0.14 |
| E3 Assertiveness | -.070 | .131 | .429 | 0.34 | -.068 | .141 | .429 | 0.32 | -.095 | .041* | .311 | 0.86 |
| E4 Activity | -.085 | .066 | .398 | 0.58 | -.057 | .222 | .527 | 0.23 | -.087 | .061 | .390 | 0.62 |
| E5 Excitement-Seeking | -.007 | .877 | .959 | 0.11 | -.048 | .302 | .632 | 0.18 | -.012 | .798 | .951 | 0.11 |
| E6 Positive Emotions | -.058 | .216 | .527 | 0.23 | -.034 | .469 | .797 | 0.14 | -.074 | .109 | .427 | 0.39 |
| Openness | | | | | | | | | | | | |
| O1 Fantasy | -.070 | .129 | .429 | 0.34 | -.063 | .178 | .515 | 0.27 | -.076 | .101 | .427 | 0.41 |
| O2 Aesthetics | -.076 | .101 | .427 | 0.41 | -.079 | .091 | .427 | 0.45 | -.118 | .011* | .149 | 2.78 |
| O3 Feelings | -.081 | .080 | .423 | 0.50 | -.062 | .183 | .515 | 0.26 | -.101 | .030* | .242 | 1.14 |
| O4 Actions | -.017 | .718 | .923 | 0.12 | -.056 | .231 | .531 | 0.22 | -.090 | .053 | .368 | 0.69 |
| O5 Ideas | -.072 | .119 | .429 | 0.36 | -.102 | .027* | .242 | 1.23 | -.135 | .004* | .126 | 7.27 |
| O6 Values | .030 | .524 | .827 | 0.13 | .002 | .967 | .984 | 0.11 | -.011 | .818 | .951 | 0.11 |
| Agreeableness | | | | | | | | | | | | |
| A1 Trust | -.058 | .214 | .527 | 0.23 | -.015 | .754 | .943 | 0.11 | -.053 | .255 | .547 | 0.21 |
| A2 Straightforwardness | -.008 | .865 | .959 | 0.11 | -.021 | .659 | .896 | 0.12 | .002 | .973 | .984 | 0.11 |
| A3 Altruism | .010 | .823 | .951 | 0.11 | -.026 | .572 | .840 | 0.13 | -.037 | .430 | .764 | 0.15 |
| A4 Compliance | .041 | .382 | .715 | 0.16 | .028 | .551 | .830 | 0.13 | .011 | .809 | .951 | 0.11 |
| A5 Modesty | .038 | .419 | .764 | 0.15 | .070 | .134 | .429 | 0.33 | .054 | .242 | .531 | 0.21 |
| A6 Tender-Mindedness | -.128 | .006* | .126 | 4.93 | -.101 | .029* | .242 | 1.14 | -.124 | .007* | .133 | 3.84 |
| Conscientiousness | | | | | | | | | | | | |
| C1 Competence | -.046 | .326 | .652 | 0.17 | -.023 | .623 | .862 | 0.12 | -.058 | .214 | .527 | 0.23 |
| C2 Order | -.007 | .884 | .959 | 0.11 | .002 | .967 | .984 | 0.11 | -.028 | .551 | .830 | 0.13 |
| C3 Dutifulness | -.030 | .518 | .827 | 0.13 | -.002 | .968 | .984 | 0.11 | -.020 | .667 | .896 | 0.12 |
| C4 Achievement Striving | -.116 | .012* | .149 | 2.51 | -.115 | .013* | .149 | 2.28 | -.132 | .005* | .126 | 5.98 |
| C5 Self-Discipline | -.031 | .506 | .827 | 0.13 | -.010 | .834 | .951 | 0.11 | -.036 | .433 | .764 | 0.15 |
| C6 Deliberation | .041 | .381 | .715 | 0.16 | .034 | .460 | .796 | 0.14 | .025 | .598 | .840 | 0.12 |

Effects of sex, age, and daytime of EEG-assessment were partialled out. FDR: False Discovery Rate according to Benjamini and Hochberg; $BF_{10}$: Bayes factor showing the likelihood ratio between the alternate and null hypothesis (1/3 beta prior width)

* $p < .05$ (two-sided nominal significance)
